# Annotation of phenotypes using ontologies: a Gold Standard for the training and evaluation of natural language processing systems

**DOI:** 10.1101/322156

**Authors:** Wasila Dahdul, Prashanti Manda, Hong Cui, James P. Balhoff, T. Alexander Dececchi, Nizar Ibrahim, Hilmar Lapp, Todd Vision, Paula M. Mabee

**Affiliations:** University of South Dakota; University of North Carolina at Greensboro; University of Arizona; University of North Carolina at Chapel Hill; University of Chicago; Duke University

**Keywords:** ontology, phenotype, annotation, anatomy, vertebrates, natural language processing

## Abstract

Natural language descriptions of organismal phenotypes - a principal object of study in biology, are abundant in biological literature. Expressing these phenotypes as logical statements using formal ontologies would enable large-scale analysis on phenotypic information from diverse systems. However, considerable human effort is required to make the semantics of phenotype descriptions amenable to machine reasoning by (a) recognizing appropriate ontological terms for entities in text and (b) stringing these terms into logical statements. Most existing Natural Language Processing tools stop at entity recognition, leaving a need for tools that can assist with both aspects of the task. The recently described Semantic CharaParser aims to meet this need. We describe the first expert-curated Gold Standard corpus for ontology-based annotation of phenotypes from the systematics literature. We use it to evaluate Semantic CharaParser’s annotations and explore differences in performance between humans and machine. We use four annotation accuracy metrics that can account for both semantically identical and similar matches. We found that machine-human consistency was significantly lower than inter-curator (human–human) consistency. Surprisingly, allowing curators access to external information that was not available to Semantic CharaParser did not significantly increase the similarity of their annotations to the Gold Standard nor have a significant effect on inter-curator consistency. We found that the similarity of machine annotations to the Gold Standard increased after new ontology terms relevant to the input text had been added. Evaluation by the original authors of the character descriptions indicated that the Gold Standard annotations came closer to representing their intended meaning than did either the curator or machine annotations. These findings point toward ways to better design of software to augment human curators, and the Gold Standard corpus will allow training and assessment of new tools to improve phenotype annotation accuracy at scale.

## 1 Introduction

Phenotype descriptions of organisms are documented across nearly all areas of biological research including biomedicine, evolution, developmental biology, and paleobiology. The vast majority of such descriptions are expressed in the scientific literature using natural language. While allowing for rich semantics, natural language descriptions can be difficult for non-experts to understand, and are opaque to machine reasoning, and thus hinder the integration of phenotypic information across different studies, taxonomic systems, and branches of biology (1).

To make phenotype descriptions more amenable to computation, model organism databases employ human curators to convert natural language phenotype descriptions into machine-readable phenotype annotations that use standard ontologies (e.g., 2, 3, 4, 5). One format used for phenotype annotations is the ontology-based Entity–Quality (EQ) representation, in which an entity represents a biological object such as an anatomical structure, space, behavior, or a biological process; a quality represents a trait or property that an entity possesses, e.g., shape, color, or size; and an optional related entity allows for binary relations such as adjacency (6, 7). The EQ format is widely used, e.g., (8), although other formal representations of phenotype descriptions have been proposed (9). Further, to create entities and qualities that adequately represent the often highly detailed phenotype descriptions, curators create complex logical expressions called ‘post-compositions’ by combining ontology terms, relations, and spatial properties in different ways. In contrast to EQ expressions with single-term entities and qualities, creating post-composed entities and qualities (Table 1) can be a complex task, due to the flexibility in logic expression and the different semantic interpretations that free-text descriptions often allow. As a result, it can be expected that EQ annotations involving post-compositions will show some variability between different curators.

**Table 1:**
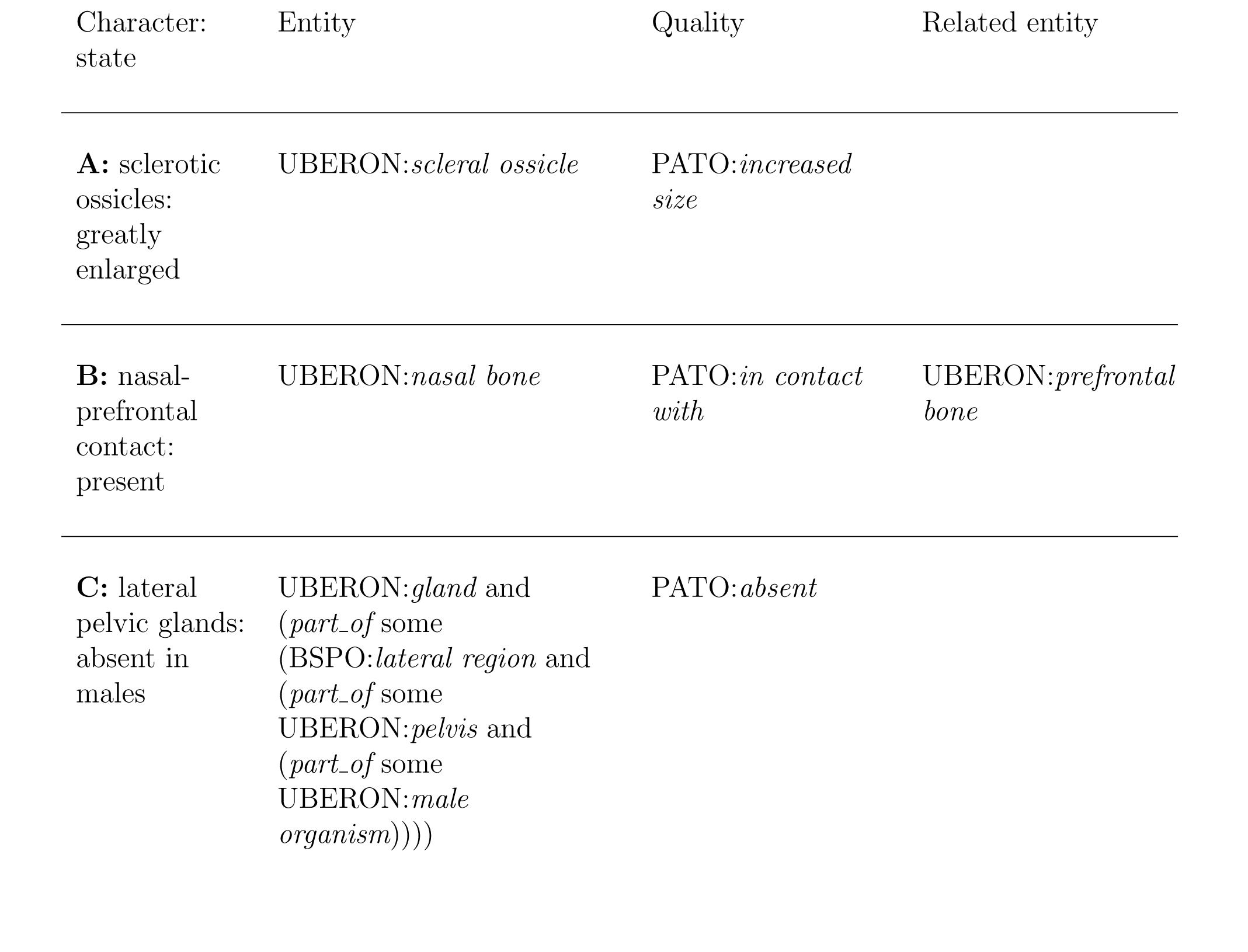
Examples of Entity–Quality (EQ) annotations of varying complexity from the present study. **A** illustrates a simple EQ annotation; **B** shows an EQ annotation in which the quality term relates two entities to each other; and **C** provides an example of an entity that does not correspond to a term in an existing ontology, but is instead a complex logical expression post-composed from multiple ontology terms.

| Character:<br>state | Entity | Quality | Related entity |
| --- | --- | --- | --- |
| **A:** sclerotic<br>ossicles:<br>greatly<br>enlarged | UBERON:*scleral ossicle* | PATO:*increased*<br>*size* |  |
| **B:** nasal-<br>prefrontal<br>contact:<br>present | UBERON:*nasal bone* | PATO:*in contact*<br>*with* | UBERON:*prefrontal*<br>*bone* |
| **C:** lateral<br>pelvic glands:<br>absent in<br>males | UBERON:*gland and*<br>*(part_of some*<br>*(BSPO:lateral region and*<br>*(part_of some*<br>*UBERON:pelvis and*<br>*(part_of some*<br>*UBERON:male*<br>*organism))))* | PATO:*absent* |  |

To best resolve the ambiguities inherent in natural language descriptions, human curators will often not only use their domain expertise, but also refer to external information for deducing the original author’s intent. Phenotype descriptions found in the literature, however, are typically in a concise format with little or no contextualizing information that would help with disambiguating the intended meaning. The difficulty of disambiguation can be exacerbated when the requisite entity and quality domain ontologies do not yet include an obviously appropriate term for a particular annotation (10). As a consequence of this and other challenges, manual curation tends to be extremely labor-intensive, and few projects have the resources to comprehensively curate the relevant literature. To help address this bottleneck, text mining and natural language processing (NLP) systems have been developed with the goal of supplementing or augmenting the work of human curators. Facilitating continuous improvement of these systems, tools, and algorithms requires means to compare different systems objectively and fairly with each other and with human curators, in particular with respect to accuracy of generated annotations. This raises several questions. One, what is the reference against which accuracy is best assessed if annotations generated for a given task show variability between different human curators? Two, how consistent is the result of machine annotation with that of a human curator? Three, to what extent is machine annotation performance limited by inherent differences between how a machine and a human expert execute a curation task? In particular, in contrast to human curators who will consult external information, a software tool will normally only use the vocabulary and domain knowledge it is initially provided with in the form of input lexicons and ontologies.

The variability among expert curators can be used to provide a baseline for the performance evaluation of automated systems. Cui et al. (11) conducted an inter-curator consistency experiment to evaluate Semantic CharaParser (SCP), a natural language processing tool designed for generating EQ annotations from phenotype descriptions in the comparative anatomy literature (specifically, from phylogenetic character matrices (12)). To our knowledge, SCP is the first semi-automatic software designed to generate EQ annotations. SCP works by parsing the original descriptions to identify entity and quality terms, matching these terms to ontology concepts, and generating logical relations and, where appropriate, post-compositions from the matched concepts based on a set of rules. In the experiment, three curators independently annotated a set of 203 characters, randomly chosen from seven publications representing extant and extinct vertebrates for a variety of anatomical systems with an emphasis on skeletal anatomy, corresponding to the curators’ domain of expertise (Table 2). In the first, or “Naïve”, round of annotation, curators were not allowed access to sources of knowledge external to the character description, including the publication from which the matrix originated. In the second, or “Knowledge” round, curators were allowed to access external sources of knowledge, such as the full publication from which the character was drawn, related literature and other online sources. The curators were given a set of initial ontologies to use for curation. The new ontology terms created during curation were added to the “Initial” ontologies to create curator-specific “Augmented” ontologies. At the end of the curation rounds, all curator-specific augmented ontologies were merged to create a final “Merged” ontology.

**Table 2:**
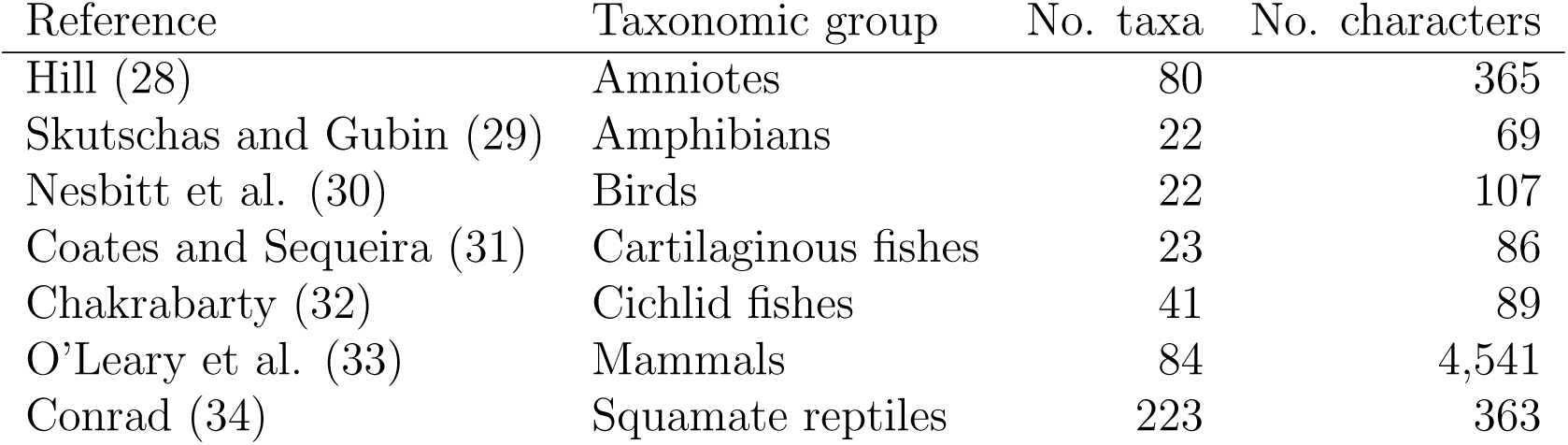
Phylogenetic studies from which characters were selected.

| Reference | Taxonomic group | No. taxa | No. characters |
| --- | --- | --- | --- |
| Hill (28) | Amniotes | 80 | 365 |
| Skutschas and Gubin (29) | Amphibians | 22 | 69 |
| Nesbitt et al. (30) | Birds | 22 | 107 |
| Coates and Sequeira (31) | Cartilaginous fishes | 23 | 86 |
| Chakrabarty (32) | Cichlid fishes | 41 | 89 |
| O’Leary et al. (33) | Mammals | 84 | 4,541 |
| Conrad (34) | Squamate reptiles | 223 | 363 |

The Cui et al. (11) study was designed such that SCP was used to annotate the same set of characters as human curators using three sets of ontologies (Initial, Augmented, and Merged) with progressively more comprehensive coverage, as described below. The primary findings were as follows. The performance of SCP was significantly lower as compared to human curators. When comparing the performance of SCP to human curators, no statistically significant differences were found between Naïve and Knowledge rounds. Inter-curator Recall and Precision were also not found to be significantly different between the Naïve and Knowledge rounds. SCP performed significantly better with Augmented versus Initial ontologies. However, there was no significant difference in performance between Augmented and Merged ontologies.

While useful, there were several limitations in the Cui et al. (11) evaluation of SCP, including the lack of a Gold Standard against which to measure its performance. Manually annotated Gold Standard datasets are high quality benchmarks for both evaluation and training of automated NLP systems e.g., (13, 14, 15). Another limitation was the use of performance measures that did not fully account for the continuum of similarity possible between semantic phenotype annotations. While these authors recognized that phenotypes annotated with parent and daughter terms in the ontology bear some partial resemblance, here we introduce semantic similarity measures that can account for any level of relationship between the terms from two phenotype annotations.

The present work describes the development of an expert-curated Gold Standard dataset of annotated phenotypes for evolutionary biology that is the best available given current constraints in semantic representation. The Gold Standard was developed for the annotation of the complex evolutionary phenotypes described in the systematics literature for the Phenoscape project (12, 16). Unlike many published gold standards for ontology annotation, which frequently focus on entity recognition, e.g., (17), the Phenoscape Gold Standard consists of EQ expressions of varying complexity. We evaluate how well the annotations of individual curators and the machine (SCP) compare to those of the Gold Standard, using four ontology-aware metrics. Two of these are traditional measures of semantic similarity (18) and two are extensions of Precision and Recall that account for partial semantic similarity. In addition, we directly assessed the quality of the Gold Standard with an author survey, in which the original domain experts were invited to rank the accuracy of a subset of the annotations from the Gold Standard, the individual human curators, and SCP.

## 2 Related Work

Gold standard corpora are collections of articles manually annotated by expert curators, and they provide a high quality comparison against which to test automated text processing systems. Funk et al. (15), for example, used the CRAFT annotation corpus (17, 19) for the evaluation of three concept annotation systems. Within the biomedical sciences, a number of Gold Standard corpora have been developed (20, 21, 22), and these focus on concept recognition. Concepts are annotated at the text string level, e.g., (17) or in some cases, annotations are attached at the whole document level, e.g., (21). Because of the effort and costs required for manual annotation, “silver standard” corpora have also been created, in which automatically generated annotations are grouped into a single corpus (23, 24). As far as we are aware, there are no published Gold Standard corpora for EQ phenotypes, and none for evolutionary phenotypes.

Inter-curator consistency has been used by several studies as a baseline against which to evaluate the performance of automated curation software (25, 26, 27). Weigers et al. measured the performance of text mining software that identifies chemical–gene interactions from the literature by comparing the output against inter-curator consistency on the same task (25). Sohngen et al. evaluated the performance of the DRENDA text-mining system, which retrieves enzyme-related information on diseases (26). Most similar to the work reported here is the study by Camon et al. (27) in which inter-curator consistency was used as a baseline to evaluate performance of text mining systems to retrieve Gene Ontology terms from literature. In their experiment, three curators co-curated 30 papers and extracted GO terms from the text. In inter-curator comparisons, GO term pairs were classified into three categories: exact matches, same lineage (terms related via subsumption relationships), and different lineage (unrelated terms). They found that curators chose exactly the same terms 39%, related terms 43%, and unrelated terms 19% of the time. Inter-curator consistency showed 94% precision and 72% recall, setting a high bar for automated text mining systems. Our approach differs in that we evaluate inter-curator consistency at the task of phenotype (EQ) annotation, and we employ metrics that can account for partial matches between annotations by taking advantage of both ontology structure and the information content from annotation frequencies.

## 3 Methods

### 3.1 Source of phenotypes

Twenty-nine phenotypic characters were randomly selected from each of seven published phylogenetic studies, yielding 203 characters and 463 character states in total (Table 2). The studies were chosen to (i) have a wide taxonomic breadth across vertebrates, (ii) include both extinct and extant taxa, and (iii) include characters from several anatomical systems (e.g., skeletal, muscular, nervous systems). These objectives were intended to reduce potential sources of systematic bias. For example, the prevailing style of character descriptions can differ depending on the taxonomic group of interest. Further, the curators had varying expertise across the vertebrate taxa. Curators had access to the full character and state descriptions, including taxonomic scope and publication source, but they—and the SCP developers—were blind to the choice of papers and the selection of characters prior to the experiment.

### 3.2 Experimental design

The common set of character states was annotated independently by three curators (W. Dahdul, T. A. Dececchi and N. Ibrahim) and by Semantic CharaParser (SCP). The curators were randomly assigned identifiers C1, C2, and C3 at the beginning of the study. Curators used Phenex software (10, 35) for manually generating annotations. The annotations are complex expressions made up of entity (E), quality (Q) and where required, a related entity (RE). The E and RE components employ Uberon (36, 37) concepts and may be post-composed with terms from multiple ontologies including Uberon, PATO (38, 39), and the Spatial Ontology (BSPO) (40) while the Q component uses PATO concepts. Curators were free to create one or multiple EQ annotations per state, and they were encouraged to annotate at a fine level of detail (41). To measure the effect of external knowledge on inter-curator consistency, two rounds of human curation were performed. In the first (“Naïve”) round, the character and character state text were the only information the curators were allowed to consult. Accessing the source publication or any external information was not permitted. This was intended to simulate the extent of information available to SCP, although curators naturally use their subject domain expertise when composing annotations. In the second (“Knowledge”) round, the curators annotated the same set of characters as in the Naïve round, but they were free to consult the full text of the source publication and to access any other external information. In total, this resulted in six sets of human-curated EQ annotations, and six augmented ontologies produced by the curators independently during the Naïve and Knowledge rounds.

Several steps were taken to promote consistency among the human curators, and between curators and SCP. First, curators developed and were trained on a set of curation guidelines for the annotation of phylogenetic characters (the Phenoscape Guide to Character Annotation (42)). These guidelines were also made available to SCP developers, and are the basis of rules according to which SCP generates EQ expressions. Second, curators took advantage of an interactive Consistency Review panel available in Phenex, which reports missing or problematic annotations, such as a relational quality used to annotate a character state without also specifying a related entity. Further, each curator had at least one year of experience with EQ annotation prior to the experiment. Note that each curator still performed their curation tasks in the experiment independently from each other, and thus there was still room for variation. For instance, for a given character state, one curator might choose to use an imperfectly matching entity term, while another might aim for a more precise representation by post-composing a new term from existing terms, and yet another might choose to add a new single term to their Initial ontology. To avoid advantaging SCP beyond an initial training dataset, SCP developers were not allowed to observe the human curation process during the experiment.

### 3.3 The Gold Standard

The Gold Standard corpus, which consists of a unique set of EQ annotations for each character state in the 203 character dataset, was created as a consensus dataset by the three curators. After completing the Knowledge round, the curators reviewed and discussed all the EQs in their three separate Knowledge round curator datasets for the purpose of developing a single Gold Standard dataset. In assembling this set of EQ annotations for the Gold Standard, the curators were not limited to choosing among the individual EQs that they had created during the experiment; instead, they were free to modify existing annotations or create entirely new ones. In cases where there was insufficient information to resolve ambiguities, the curators consulted additional published literature and other online resources. In some cases, they also contacted domain experts to clarify terminology or anatomy. Once all three curators were in agreement, they used the Phenex curation software to create the Gold Standard EQ annotations for the final Gold Standard dataset.

In the course of developing the Gold Standard, the curators updated the Phenoscape Guide to Character Annotation (42) with previously missing character categories, examples, and EQ conventions. Each phenotype in the Gold Standard was categorized according to one or more of the character types from this Guide.

### 3.4 Ontologies

The human curators and SCP were provided with the same initial set of ontologies: the Uberon anatomy ontology (version phenoscape-ext/2013-03-15, (36, 37)), the Spatial Ontology (BSPO) (release 2013-05-17, (40)), and the Phenotype and Trait Ontology (PATO) (release 2013-06-03, (39)).

In both the Naïve and Knowledge rounds, each curator was free to provisionally add terms that they deemed missing from any of the Initial ontologies, resulting in Augmented ontologies that differed from their Initial versions. New term requests were added as provisional terms by using the Ontology Request Broker in Phenex (10), which provides an interface to the BioPortal’s provisional term API (43). Ontology curators can subsequently resolve these requests as mistakenly overlooked existing terms, new synonyms to existing terms, or *bona fide* new terms. At the end of the experiment, there were six sets of Augmented ontologies, one from each curator in each round (Table 3). These were subsequently combined to produce a Merged set of ontologies for which redundant classes were manually reconciled. To test the effect of ontology coverage on automated EQ annotation, SCP was run with the Initial ontology, the Augmented ontologies, and the final Merged ontology. The results in each case were compared to those obtained by the human curators, as reported in Cui et al. (11).

**Table 3:** Augmentation of entity (UBERON), quality (PATO), and spatial (BSPO) ontologies by the three curators in both rounds of curation (Naïve and Knowledge). The final Merged ontology includes the reconciled set of terms from all six Augmented ontologies.

| Curation round | Human curator | Terms added to: |  |  |
| --- | --- | --- | --- | --- |
|  |  | UBERON | PATO | BSPO |
| Naïve | C1 | 109 | 70 | 3 |
|  | C2 | 49 | 32 | 0 |
|  | C3 | 89 | 23 | 2 |
| Knowledge | C1 | 129 | 74 | 3 |
|  | C2 | 72 | 52 | 0 |
|  | C3 | 108 | 35 | 3 |
| Merged |  | 199 | 127 | 7 |

### 3.5 Measuring similarity between annotation sources

When different ontology terms are chosen to annotate a given character state, the selected terms may nonetheless be semantically similar. Thus, it is desirable to use measures of annotation similarity that allow for varying degrees of relatedness using the background ontology and annotation corpus (18). Here, we use four measures, two of which are semantic similarity metrics with a history of usage in the literature, and two of which are modifications of the traditional measures of Precision and Recall that account for different but semantically similar annotations. All four measures can be applied to both full EQ annotations and to comparisons among entity terms alone.

Semantic similarity measures between annotation sources (e.g., different curators) were aggregated at the level of the individual character state, and across all character states (Figure 1). Aggregation of pairwise (EQ to EQ) annotations by character state is necessary because a curator may generate more than one EQ annotation for a given character state. This is illustrated by Figure 1 where Curator A generated three EQs and Curator B generated two EQs for State *i*. To measure the overall similarity between two annotation sources (e.g., Curator A to Curator B in Figure 1, top), first, we compute similarity between corresponding character state pairs. For each character state pair, we take the best match (maximum score) among all pairwise comparisons between EQs for the same character state (Maximum Character State Similarity in Figure 1). Second, we take the mean of the similarity scores for all character state pairs (Mean Curator Similarity in Figure 1, bottom).

**Figure 1:**
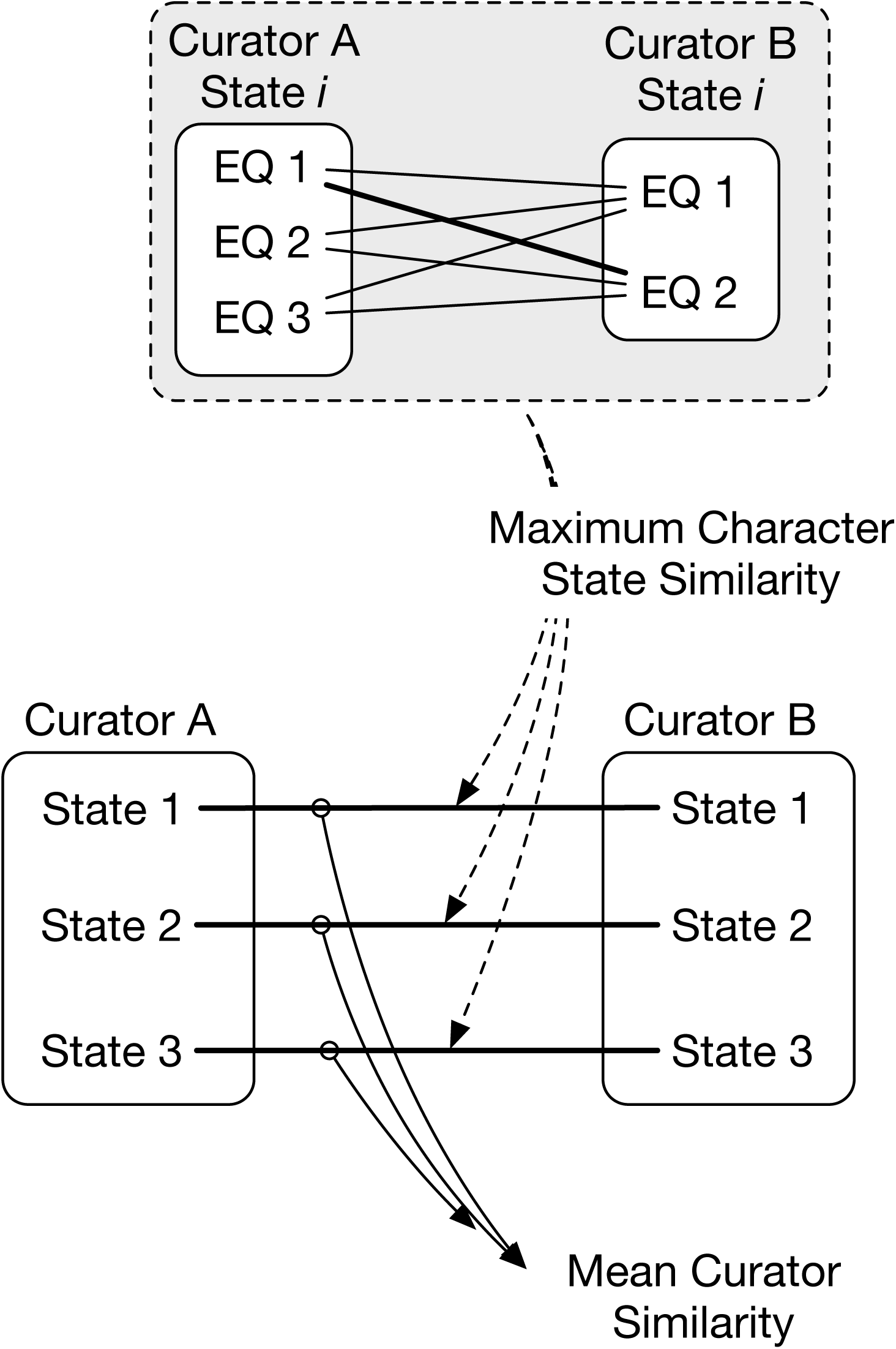
Similarity of annotations between two curators is calculated across multiple character states (e.g., states 1–3, bottom). First, the maximum character state similarity is calculated at the level of a single character state, and is the best match (maximum score) in pairwise comparisons across that state’s EQ annotations. Mean curator similarity is then calculated as the mean of the maximum similarities across all character state pairs.

#### 3.5.1 Generating subsumers for EQ annotations

We treat each EQ annotation as a node in an *ad hoc* EQ ontology. Creating the complete cross-product of the component ontologies would necessarily include all possible subsumers but would be prohibitive. As a memory saving measure, we developed a computationally efficient approach to identify subsumers for EQ annotations on an *ad hoc* basis, as follows.

A comprehensive ontology was created by taking the union of Uberon, PATO and BSPO ontologies using the *–merge-support-ontologies* command in the owltools software (https://github.com/owlcollab/owltools). In order to enable reasoning on additional dimensions (e.g., *part of*) in post-compositions while identifying subsumers, we added additional classes to the comprehensive ontology. For every concept *U* in the Uberon ontology and every object property *OP* used in post-compositions, a class of the form “*OP* some *U*” was added to the comprehensive ontology.

First, every EQ annotation is split into individual E, Q, and optionally, RE components (Figure 2, Step 1). Simultaneously, the EQ annotation is transformed into an OWL class expression of the form “Q and inheres in some E and towards some RE” (Figure 2, Step 1). Next, superclasses of these individual components and the class expression are retrieved using the ELK reasoner on the comprehensive ontology (Figure 2, Step 2). Individual E, Q, RE superclasses are combined to create superclasses of the form E-Q-RE. The combined class expression and combinatorial E-Q-RE superclasses form the subsumers of an EQ annotation (Figure 2, Step 3). While it is possible that additional subsumers could be found in the case that a class in another part of the hierarchy has a logical definition that matches an EQ expression, it is unlikely for these ontologies because subsuming quality terms in the PATO ontology do not have logical definitions which make use of Uberon entities.

**Figure 2:**
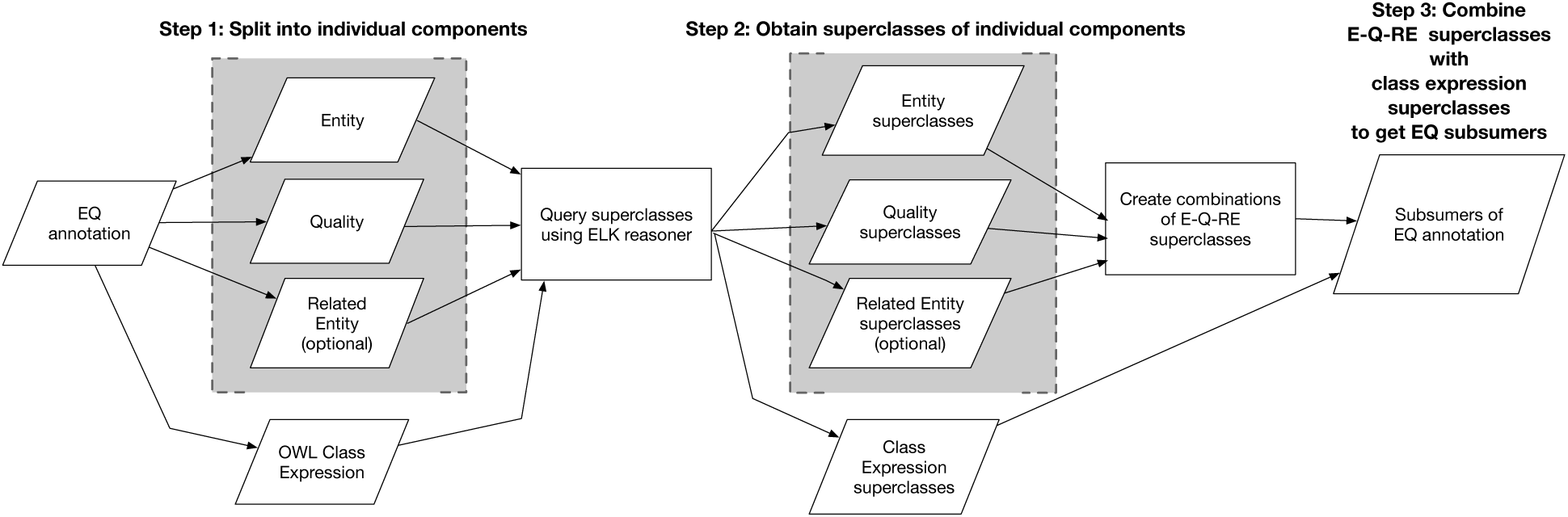
EQ annotations are split into Entity (E), Quality (Q), and Related Entity (RE) components, and also, transformed into an OWL class expression. Superclasses of E, Q, RE, and the class expression are queried via ELK. E, Q, RE superclasses are combined in the form E-Q-RE. These E-Q-RE superclasses along with the class expression’s superclasses form the subsumers of the EQ annotation for computation of semantic similarity.

#### 3.5.2 Jaccard Similarity

The Jaccard Similarity (*J*_sim_) between nodes *N*_1_ and *N*_2_ in an ontology graph is defined as the ratio of the number of nodes in the intersection of their subsumers over the number of nodes in the union of their subsumers (44):

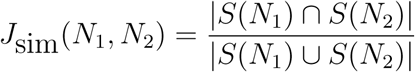

where *S*(*N_i_*) is the set of nodes that subsume *N_i_*. *J*_sim_ measures the distance between two EQs based on the class structure of the ontology. The range of *J*_sim_ = [0,1]. *J*_sim_ = 1 when the two EQs being compared are the same and *J*_sim_ = 0 when they have no common subsumers.

#### 3.5.3 Information Content

*J*_sim_ measures the ontology graph distance between two nodes, and thus necessarily ignores differences in semantic specificity between parent and child terms in different areas of the ontology graph. Information Content (*IC*) is used to capture the specificity of the annotations. The Information Content *I* of a node *N_j_* in an ontology is defined as the proportion of annotations to *N_j_* and all nodes subsumed by *N_j_* in an annotation corpus (45). Let *q* be the number of nodes in the ontology. Define *f(N*) to be the number of annotations directly to *N_j_* and *g*(*N*) to be the sum of annotations to *N_j_* and to all nodes subsumed by *N_j_*:

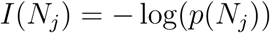

where

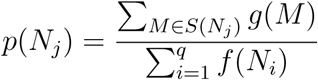

The *I* of two nodes is defined as the *I* of the Least Common Subsumer (LCS) of the two nodes. If there are multiple LCSs, the node with the highest *I* is used (44). *I* has a minimum of zero at the root and a maximum that is dependent on the size of the corpus

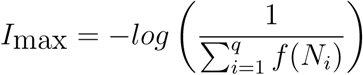

To obtain a normalized score *I_n_* in the range of [0,1], we use *I_n_* = *I/I_max_*. In our analysis, the corpus for measurement of *I_n_* includes all human annotations from both annotation rounds and the annotations from SCP.

#### 3.5.4 Partial Precision and Partial Recall

Precision and Recall are commonly used to evaluate the performance of information retrieval systems. Traditionally, these two measures do not attempt to account for imperfect matches; information is either retrieved or it is not. For ontology-based annotations, partial information retrieval is possible because the information to be retrieved is the semantics of the annotated text, rather than a particular term. To account for this, here we use two metrics, Partial Precision (*PP*) and Partial Recall (*PR*), to measure the success of semantic information retrieval by a test curator (*C_T_*) relative to a reference curator (*C_R_*), where a curator can be understood as either human or software. While other variants of semantic precision and recall are used in the literature (46, 47), the measures we use here specifically use semantic similarity, in this case *J*_sim_, to quantify partial matches between annotations. In contrast to our approach, (46) and (47) compute semantic precision and recall by examining the superclass sets of two annotations. Depending on the overlap among these sets, each superclass is classified as a true positive, false positive, or false negative. These counts are then used to compute semantic precision and recall.

PP measures the proportion of the semantics annotated by *C_R_* that are retrieved by *C_T_* relative to the number of *C_T_* annotations. *PR*, on the other hand, measures the proportion of semantics that are retrieved by *C_T_* relative to the number of *C_R_* annotations. Thus, both *PP* and *PR* have a range of [0,1]. *PP* will decrease due to extra annotations by *C_T_* that are dissimilar from those in *C_R_*, while PR will decrease due to extra annotations in *C_R_* that are lacking from *C_T_*. Both use *J*_sim_ to measure semantic similarity and are computed at the character-state level rather than the individual EQ annotation level. Using *C_R_* and *C_T_* as an example, they are calculated as:

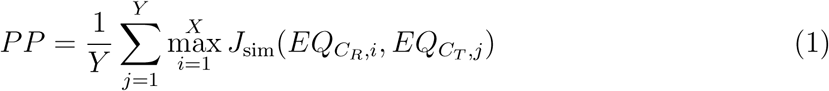

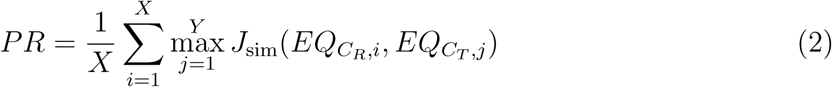

where *i* = 1..*X* indexes the EQs from *C_R_* and *j* = 1..*Y* indexes the EQs from *C_T_*.

### 3.6 Author assessment of Gold Standard, curator, and machine annotations

To assess how close EQ annotations created by the different sources came to the intent of the authors of the seven studies from which the characters were drawn, an author from each was invited to evaluate the relative performance of the annotation sources. Using SurveyMonkey (www.surveymonkey.com), we presented one author from each study with ten randomly selected character states derived from their publication and asked them to rank the five different annotation sources (C1, C2, C3, SCP, GS) for each state [Section 1, Supplementary Materials].

Authors were given background material at the beginning of the survey describing the EQ method of character annotation. Authors were then asked to rank annotations in order of preference, with the annotation that best represented the meaning of the character state ranked first. Annotations were presented in random order, and the source of each annotation could not be tracked by the author. All of the EQ annotations for each character state generated by a particular annotation source were presented to the authors.

We used two statistics to test for differences among author preferences for the different annotation sources (48). Anderson’s statistic, *A*, was used to test whether the overall distribution of ranks was different in the observed (*O*) data than expected (*X*):

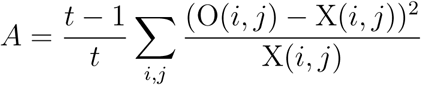

where the expected number of observations *X*(*i*, *j*) = *n/t* for factor *i* assigned rank *j* and number of observations *n*. *A* was tested against a *χ*^2^ distribution for significance with (*t* − 1)^2^ degrees of freedom. The null hypothesis is that all author preferences for all annotation sources will be equally frequent.

Friedman’s statistic, *F*, was used to test if the mean ranks of the different annotation sources differed from chance:

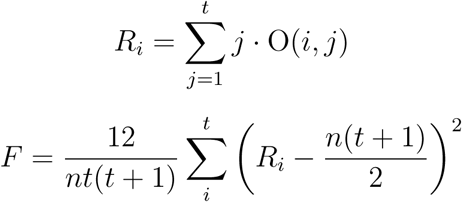

where *t* = 5 is the number of annotation sources, *i* = 1..*t* is the annotation source, *j* = 1..*t* is the number of ranks that can be assigned to an annotation, obs(*i, j*) is the number of times rank *j* was assigned to factor *i*, and *n* is the number of observations, as before. *F* was tested against a χ^2^ distribution for significance with *t* − 1 = 4 degrees of freedom.

## 4 Results

### 4.1 Datasets and source code

The Gold Standard corpus is available in NeXML (49) (`Gold_Standard-final.xml`) and spreadsheet formats (Excel: `GS-categories.xls`; tab-delimited: `GS-categories.tsv`). The files include the full-text character and character state descriptions, the source study, and the associated EQ phenotypes. The spreadsheet format also contains references for each phenotype to the character categories from the Phenoscape Guide to Character Annotation (42). The corpus in the different formats, as well as the ontologies and annotations generated in its production, have been archived at Zenodo (https://doi.org/10.5281/zenodo.1217594). The source code for the analysis of inter-curator and SCP consistency based on semantic similarity metrics, as well as the data and ontologies used as input, have been archived separately, also at Zenodo (https://doi.org/10.5281/zenodo.1218010).

Semantic CharaParser is available in source code from GitHub (https://github.com/phenoscape/phenoscape-nlp/) under the MIT license. The version used for this paper is the 0.1.0-goldstandard release (https://github.com/phenoscape/phenoscape-nlp/releases/tag/v0.1.0-goldstandard), which is also archived at Zenodo (https://doi.org/10.5281/zenodo.1246698).

### 4.2 Gold Standard

The Gold Standard dataset consists of 617 EQ phenotypes annotated for 203 characters and 463 character states. In total, these phenotypes are composed of 1,096 anatomical terms (312 unique concepts) from Uberon, 698 quality terms (147 unique) from PATO, and 148 spatial terms (30 unique) from BSPO. The dataset contains 339 post-composed terms (277 anatomical and 62 quality terms) created by relating existing terms from the same or different ontologies.

New anatomy and quality terms were required for the completion of the Gold Standard annotations. From the full set of terms individually created by the curators during the experiment (Table 3), a total of 111 anatomical terms and 12 synonyms, and 20 quality terms and two synonyms, were added to the public versions of Uberon and PATO, respectively. The remaining subset of terms created by curators in the Merged ontology were not added to the public ontology versions either because a different term was chosen for the GS annotation of a particular character, or the term was determined to be invalid after discussion among curators.

Using *J*_sim_ and *I_n_* (see Section 3.5) to measure semantic similarity between the four individual annotation sources (C1, C2, C3, SCP) and the Gold Standard, we examined (i) whether the human annotations (C1, C2, C3) showed an increase in similarity to the Gold Standard between the Naïve and Knowledge rounds and (ii) whether the machine annotations (SCP) showed an increase in similarity to the Gold Standard as ontologies progressed from the Initial, to Augmented, and to the final Merged version.

Figure 3 shows similarity (as measured by *PP*, *PR*, *J*_sim_, and *I_n_*) between annotations derived from the curators and the Gold Standard in Naïve and Knowledge curation rounds. Based on two sided, paired Wilcoxon signed rank tests, *PR* and *J*_sim_ significantly differed for C1 (*PR*: *p* =1.10 × 10^−12^, *J*_sim_: *p* = 2.06 × 10^−10^) and C2 (*PR*: *p* = 8.49 × 10^−5^, *J*_sim_: *P* = 0.0002), *PP* significantly differed for C1 (*p* = 1.24 × 10^−10^), while *I_n_* significantly differed for C1 (*p* = 2.15 × 10^−11^) between the Naïve and Knowledge rounds.

**Figure 3:**
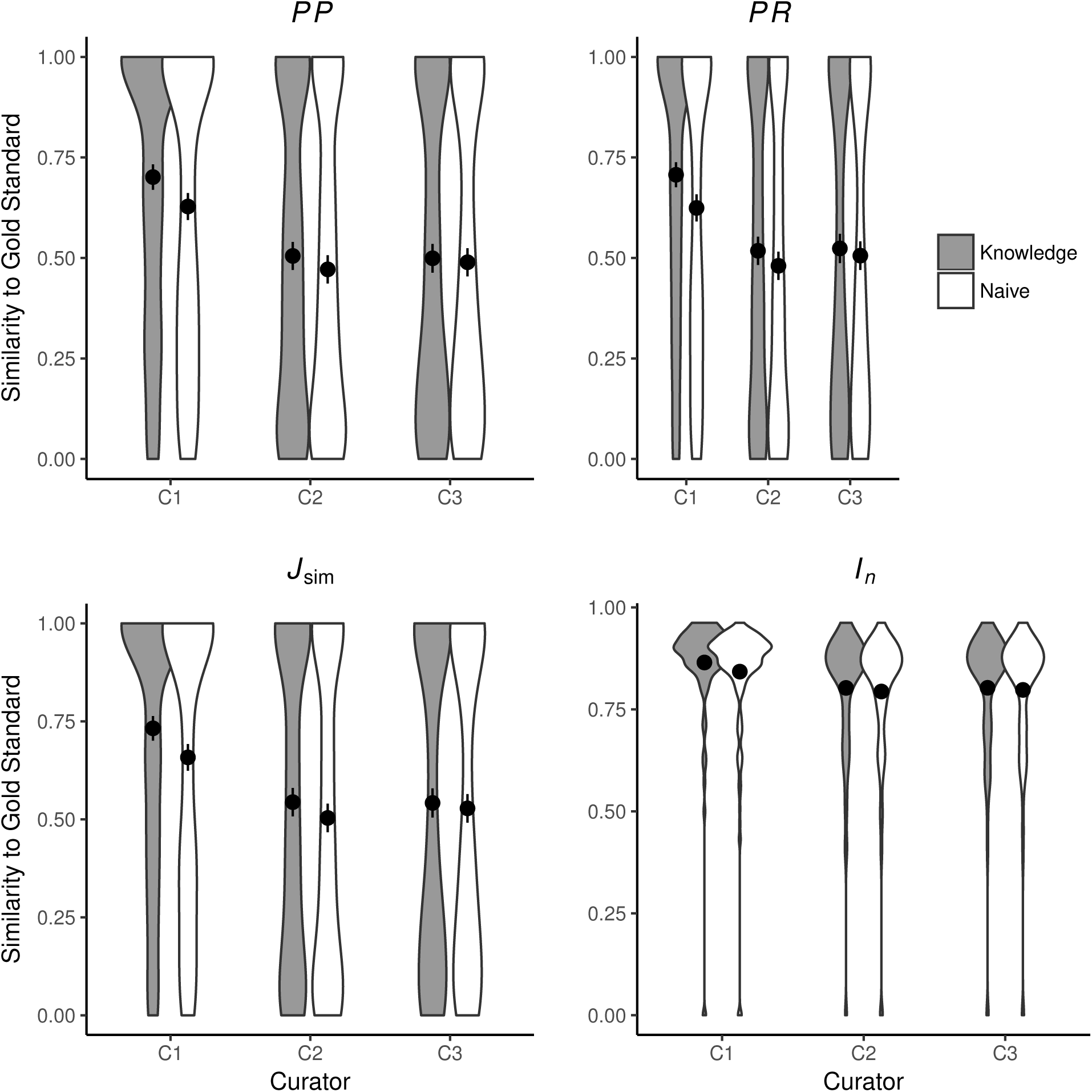
Similarity of human annotations to the Gold Standard in Naïve and Knowledge rounds. Shown are means across all 463 character states. Error bars represent two standard errors of the mean. Curators C1 (as per *PP*, *PR*, *J*_sim_, and *I_n_*) and C2 (as per *PR*, *J*_sim_) were significantly closer to the Gold Standard in the Knowledge round as compared to the Naïve round. Detailed results are shown in Supplementary Materials, Table 1

Similarity of SCP annotations to the Gold Standard increased (26% average improvement across the four metrics) after new ontology terms had been added by human curators (detailed results are in Supplementary Materials, Table 2). The majority of statistics were significantly affected between the use of the Augmented and final Merged ontologies in both annotation rounds (Figure 4) with a few exceptions. *PP* and *J*_sim_ were not affected for C1 in the Knowledge round while *PR* was not affected in both rounds for C2. For C3, *J*_sim_, *PP* in the Knowledge round and *PR* in Naïve round were not significantly affected. *p*-values for individual comparisons are shown in Supplementary Materials, Table 2.

**Figure 4:**
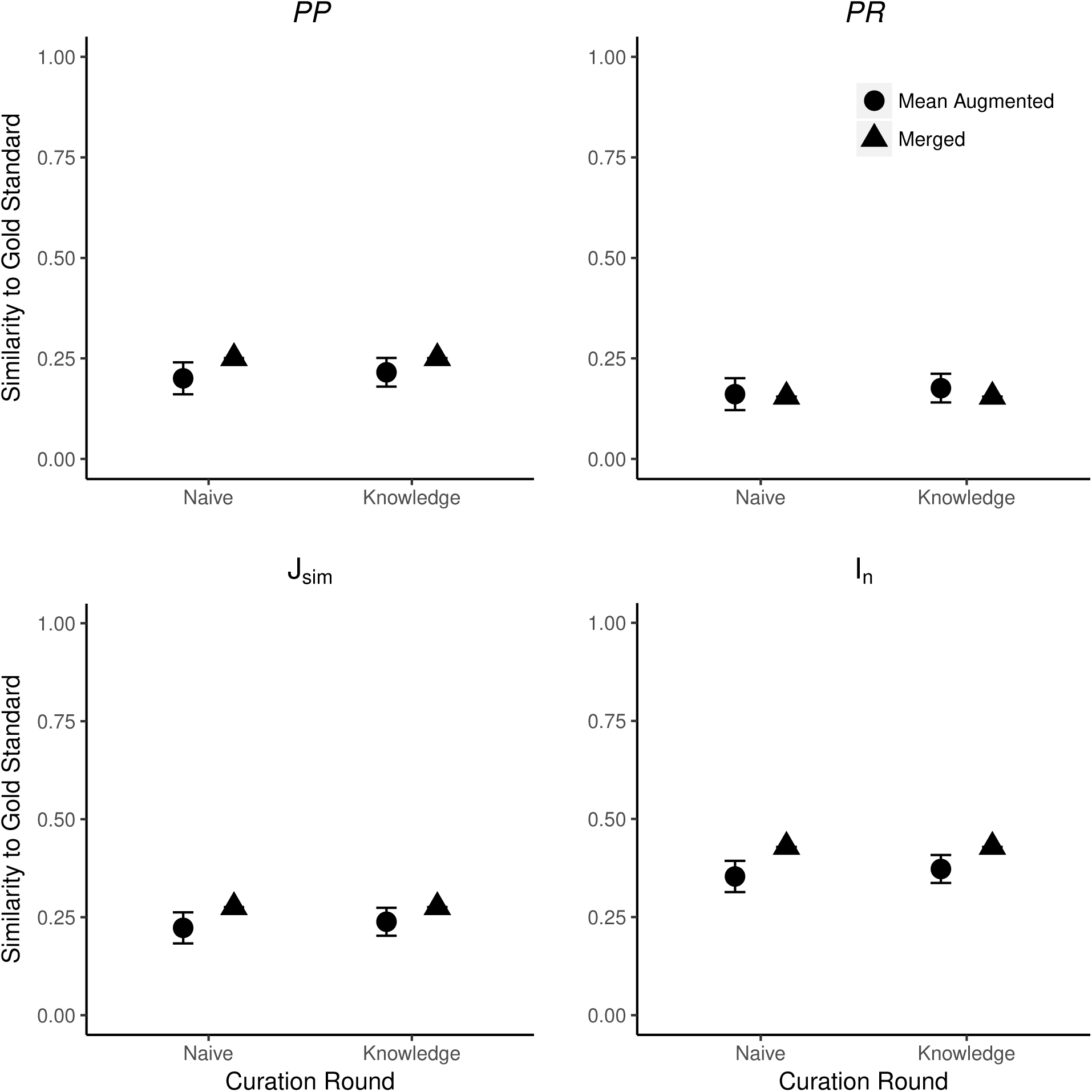
Effect of ontology completeness on SCP performance as measured by similarity to the Gold Standard. ‘Mean Augmented’ is the mean of similarity scores from the three curator augmented ontologies; error bars show two standard errors of the mean. Significant differences in similarity between SCP and the Gold Standard were found for the majority of statistics across the two rounds. Detailed results are shown in Supplementary Materials, Table 2 29% were more complex, 33% were less complex, and 38% retained the same complexity in the Knowledge round.

### 4.3 Consistency among human curators

We computed consistency among curators for the EQ annotations generated for each character state. Figure 5 shows the mean inter-curator consistency scores across three pairwise comparisons in the Naïve and Knowledge rounds respectively for Partial Precision (PP), Partial Recall (PR), *J*_sim_, and *I_n_*. The differences between Naïve and Knowledge rounds are not statistically significant (two sided, paired Wilcoxon signed rank tests, *n* = 463, *p* > 0.05 for all comparisons). These results echo those reported by Cui et al. (11) for the same experiment but reflect statistics that account for ontology structure or annotation density.

**Figure 5:**
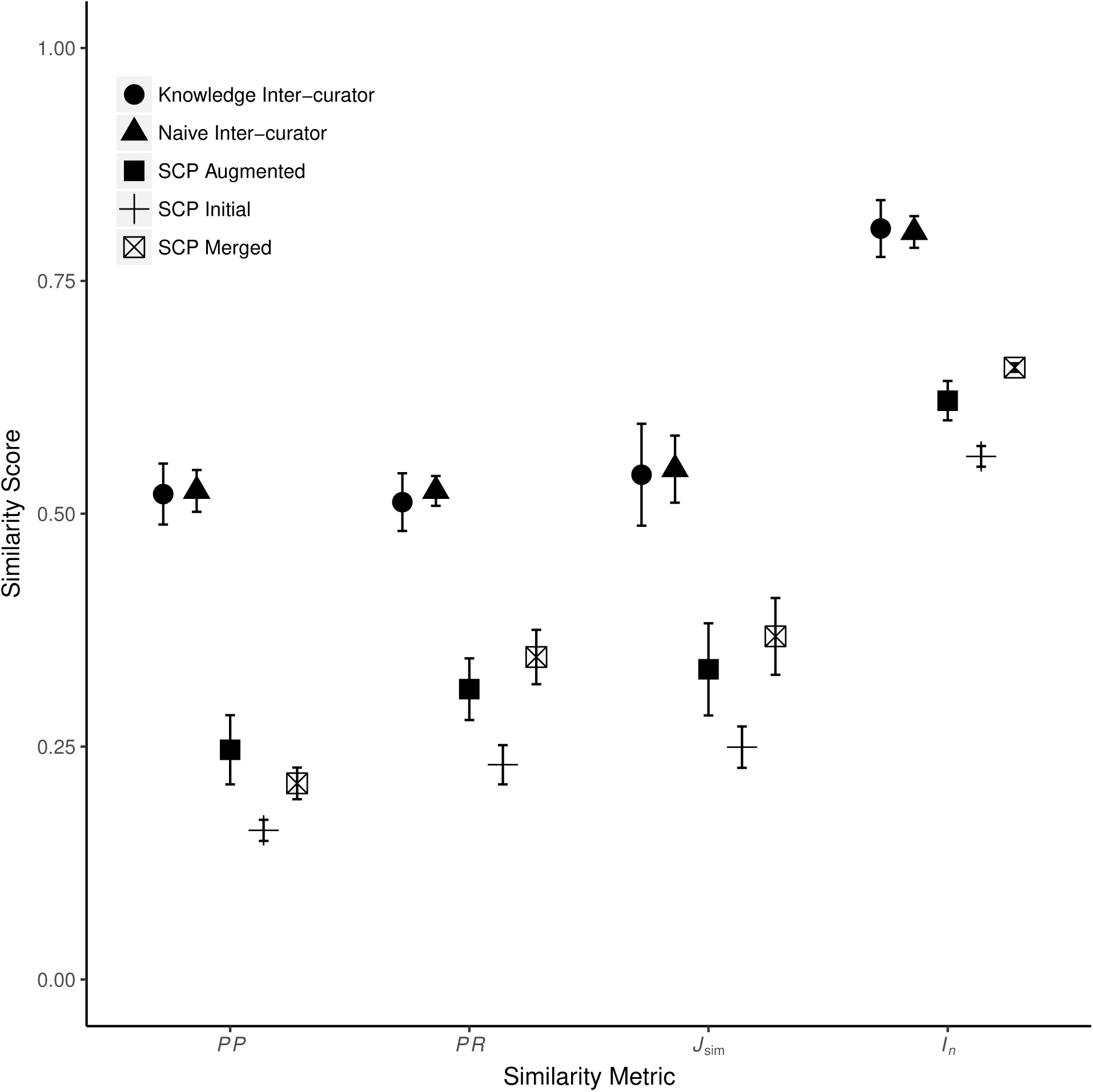
Mean inter-curator consistency and mean similarity between human and machine (SCP) generated annotations. Error bars show two standard errors of the mean. Inter-curator consistency results are shown for both the Naïve and Knowledge annotation rounds. SCP runs used either the Initial, C1, C2, or C3 Augmented, or the Merged ontologies. Only SCP similarity to human-generated annotations from the Knowledge round are shown. Consistency between SCP annotations to human annotations was significantly lower than human inter-curator consistency. Across all metrics, SCP annotation similarity to human annotations increased significantly between the use of Initial to Augmented ontologies and again from Augmented to the Merged ontology except for *PP* (decreased from Augmented to Merged). Detailed results are in Supplementary Materials, Tables 3, 4

To evaluate whether the absence of a difference in inter-curator consistency between the Naïve and Knowledge rounds was because curators made mostly the same annotations in both rounds, Cui et al. (11) examined the changes in EQ annotations. They found that curators created substantially different EQ annotations in the Knowledge round as compared to the Naïve round. Each curator changed EQ annotations between these rounds for more than 50% of character states. Among the EQs that were different between the two rounds,

Due to the lack of significant differences between inter-curator consistency in Naïve and Knowledge rounds (Figure 5), we only report curator results for the Knowledge round in subsequent sections.

### 4.4 Human–machine consistency

Using the same metrics as above, we compared the human-generated annotations to those generated by SCP. To evaluate the effect of the completeness of ontologies on SCP performance, we ran SCP separately with the Initial ontology, each of the three (C1, C2, or C3) Augmented ontologies, and the Merged ontology. Approximately 15–20% of character state annotations made by SCP using the different ontologies contained incomplete EQs. Incomplete EQs refer to those statements that are only partially matched to ontology terms, e.g., either E or Q terms are matched. In case of post-compositions, some parts needed in the composition are not matched to an ontology term. Human–machine comparisons involving character states with incomplete EQs were awarded a 0 similarity score.

We found that machine-human consistency was significantly lower than inter-curator consistency by an average of 35% across the four metrics (detailed results are in Supplementary Materials, Tables 3, 4). The overall averages for the four scores in the human–machine comparison (unfilled square markers in Figure 5) are substantially lower than the averages for the comparisons among the human curators (circle markers in Figure 5). These comparisons are statistically significant for all four metrics (two sided, paired Wilcoxon signed rank test: *PP*: *p* = 1.82 × 10^−13^; *PR*: *p* = 3.36 × 10^−43^; *J*_sim_: *p* = 7.78 × 10^−18^, *I_n_*: *p* = 9.83 × 10^−32^).

#### 4.4.1 Effect of ontology completeness on SCP-human consistency

Figure 5 shows the resulting *PP*, *PR*, *J*_sim_, and *I_n_* scores comparing SCP annotations generated with the Initial, Merged, or Augmented ontologies (plus, unfilled square, and filled square markers, respectively) to annotations from the human Knowledge round (as noted above, no statistically significant differences were found in SCP similarity to human annotations between the Naïve versus Knowledge rounds). However, almost universally, the scores among the similarity metrics increased as the ontologies progressed from Initial to Augmented and then from Augmented to Merged. The one exception is Partial Precision, which declined from the Augmented to the Merged ontology. All these increases, and the one decrease, were found to be statistically significant with two-sided paired Wilcoxon rank sum tests at the Bonferonni-corrected threshold of α = 0.0008 (Table 5).

### 4.5 Author evaluation

We received responses to survey requests from six of the seven authors of the seven source studies (Table 2). Of the six completed surveys, 3 authors evaluated (ranked) phenotypes for all 10 characters; 1 author ranked 9 characters; and 2 authors ranked 8 characters. Table 4 reports the mean rank assigned to each curation source. The overall distribution of ranks differed significantly among the curation sources (Friedman’s statistic, *p* = 0.00114) and there were significant differences among the mean ranks of each (Anderson’s statistic, *p* = 0.00133). The GS had the lowest mean rank among the annotation sources, and authors ranked the GS annotations first for 21 out of 55 characters, indicating that the GS came closest to the meaning of the original authors more frequently than others. SCP had the highest mean rank, indicating that the machine annotations were farthest away from the original authors’ intent more frequently than the individual human curators or the GS.

**Table 4:** Evaluation of annotations by original authors. Authors ranked the annotations from the Gold Standard, the three human curators (C1, C2 and C3) and Semantic Charaparser (SCP). A lower value corresponds to an annotation deemed to be more accurate or precise.

| Annotation source | Mean rank |
| --- | --- |
| Gold Standard | 2.55 |
| C1 | 2.62 |
| C2 | 3.02 |
| C3 | 3.15 |
| SCP | 3.67 |

**Table 5:** Comparison of Semantic CharaParser annotations using Initial, Augmented, and Merged ontologies to measure the effect of ontology completeness on SCP-human consistency. Shown are *p*-values from two-sided paired Wilcoxon rank sum tests.

| Comparison | $PP$ | $PR$ | $J_{\text{sim}}$ | $I_n$ |
| --- | --- | --- | --- | --- |
| Initial vs. Augmented ontologies | $9.45 \times 10^{-46}$ | $7.98 \times 10^{-39}$ | $1.67 \times 10^{-19}$ | $1.43 \times 10^{-14}$ |
| Augmented vs. Merged ontologies | $1.71 \times 10^{-15}$ | $7.26 \times 10^{-23}$ | $3.02 \times 10^{-16}$ | $8.35 \times 10^{-16}$ |

## 5 Discussion

### 5.1 Gold Standard

Phenotype curation is typically done manually, without significant assistance from machines. It is difficult and time-consuming, and across a wide variety of fields, from agriculture to medicine, it has been found not to scale to the size of the task at hand (50, 51). Developing effective machine-based methods to aid in this task, however, requires standards against which to measure machine performance. The corpus of annotations developed here as a Gold Standard is the result of a methodical, multi-step process. Beginning with the choice of seven papers in the field of phylogenetic systematics that represent phenotypic diversity across extinct and extant vertebrates, a set of 203 characters (463 states) were randomly selected. Three experienced curators with training and experience in EQ annotation and research backgrounds in vertebrate anatomy and phylogenetics independently annotated the characters while simultaneously augmenting the initial ontologies. After merging their individual augmented ontologies, the three curators then discussed their annotations for each character state, and in some cases referenced external knowledge and contacted domain experts to clarify concepts, to develop consensus annotations. We then turned to the researchers who conceived of and described the original character states to assess the consensus annotations in relation to the machine-generated and individual curator annotations. Their judgment that the consensus annotations were on average closest in meaning to their original representation in free text validates use of the consensus annotations as a Gold Standard.

The Gold Standard presented here is the first of its type for evaluation of progress in machine learning of EQ phenotypes. It differs in a number of other ways from previously published Gold Standard corpora in the biomedical sciences. Rather than ensuring that every concept in the text of a character state is tagged with an ontology term (as is the case for a concept-based Gold Standard, such as CRAFT (17)), we focused on generating EQ annotations that best represent the anatomical variation described in a character. Thus, in some cases, the EQ or EQs chosen for a particular character state may not include ontology terms in one-to-one correspondence with concepts described in the character. For example, the character state “parietal, entocarotid fossa, absent” was represented in a single EQ as E: *‘entocarotid fossa’*, Q: *‘absent’.* Parietal was not annotated because entocarotid fossa is the focus of the character, not the structure (parietal) that it is a part of. In addition, the domain knowledge that entocarotid fossa is part of the parietal is encoded in the Uberon anatomy ontology.

Similarly, in some cases, character states describing the presence of a structure are not annotated directly in the Gold Standard. This is because presence can be inferred using machine reasoning on annotations to different attributes (e.g., shape) of the structure (52). In the following character state, for example, “Hemipenis, horns: present, multi-cusped” (34), the annotation in the Gold Standard consists of a single EQ phenotype: E: *‘horn of hemipenis’*, Q: *‘multicuspidate’.* The presence of *‘horn of hemipenis’* is inferred by the assertion describing its shape and did not require a separate EQ annotation.

In other cases, “coarse” level annotations were used that did not include every concept in the character state due to limited expressivity in the EQ formalism. For example, take the character “Quadrate, proximal portion, lateral condyle separated from the medial condyle by a deep but narrow furrow”. This relates three entities (lateral condyle of quadrate, medial condyle of quadrate, furrow), which cannot be expressed using the current EQ template model in Phenex: (30). Instead, this character state was annotated coarsely as: E: *‘lateral condyle of quadrate’*, Q: *‘position’*, RE: *‘medial condyle of quadrate’*

More complex annotations can be made using a less restrictive annotation tool (e.g., Protégé) rather than the EQ templates available in Phenex. However, allowing increased complexity when annotating in EQ format is likely to increase inter-curator variability. Pre-composed ontologies, i.e., phenotype ontologies, such as used by the HPO (53), could, however, potentially decrease inter-curator variability because curators would be more likely to choose among existing annotations rather than requesting a new one. Curators would also be aided by having access to existing, vetted annotations when creating new ones. Finally, providing additional context for character descriptions, such as specimen illustrations or images, could greatly aid curators in capturing the original intent of a character. Although most publications do include illustrations or images for some characters, rarely is this done for all characters in a matrix.

Finally, in some cases the Gold Standard annotations did not fully represent the knowledge (explicit or implicit) of a character due to limitations in the expressivity of OWL. For example, in the character: “height of the vertebral centrum relative to length of the neural spine”, size is implicitly compared between two structures in the same individual. However, such within-individual comparison cannot be fully represented using an OWL class expression (54).

### 5.2 Inter-curator variation

The goal of evaluating the performance of automated curation tools is to engineer and improve machine-based curation to assist human curation as effectively as possible. Phenotype curation relies on deep domain and ontology knowledge as well as on expert judgement. Semantics in character descriptions can be variably interpreted, creating an inherent inter-curator variability. Thus, to judge the performance of automated curation tools against humans, it is important to first understand the level of variation between human curators as well as the sources of that variation.

As expected, we found considerable variation among human curators in our experiments. We observed that human curators achieved on average 54% of the maximum possible consistency as measured by *J*_sim_, and 80% as measured by *I_n_* (Figure 5). This sets a ceiling for the maximum performance of a computational system if we assume that the human variability is primarily a consequence of the inherent ambiguity in how best to capture the semantics of the phenotype statement given the available ontologies.

Much of the observed inter-curator variation could be assigned to a few general types of sources:

- Curators choose different but related terms. For example, terms may be related through subsumption (e.g., *‘circular’* and *‘subcircular’* in PATO) or sibling relationships (e.g., PATO: *‘unfused from’* and *‘separated from*’)
- Curators make differing decisions about how to post-compose entities. For example the entity for the character *“lateral pelvic glands, absent in males*” was composed differently by the three curators as *“gland and (part-of some (lateral region and part_of some pelvis))”, “lateral pelvic gland and (part_of some male organism)”*, and *“male organism and (hasjpart some (pelvic glands and in_lateral_side_of some multi-cellular organism))”.*
- Curators differ in how they composed an EQ even when choosing the same ontology terms. For example, two differently composed annotations for the character *“pelvic plate semicircular, present”* were E: *pelvic plate and (bearer_of some semicircular)* + Q: *present* and E: *pelvic plate + Q: semicircular.*
- Curators differ in how they added needed terms to the ontologies. For example, in the phenotype *“dermal sculpture on skull-roof weak“*, one curator created a new term *“surface sculpting”* and post-composed the entity *“surface sculpting and (part_of some dermatocranium)”* as the ontological translation of the entity because *“dermal sculpture“* did not exist in the Uberon anatomy ontology. Another curator used PATO: *‘sculpted surface’* to create a post-composed entity term *“dermatocranium and (bearer_of some sculpted surface)”* to represent the same entity.

### 5.3 Human–machine variation

SCP achieved, on average, 37% and 66% consistency with human curators using the most comprehensive (merged) ontology, as measured by *J*_sim_ and *I_n_*, respectively (Figure 5). This shows that the performance of SCP is significantly lower as compared to human inter-curator performance.

### 5.4 Usefulness of semantic similarity for partial matches

One of the major sources of annotation variation in either human or machine curators stems from choosing terms that are related to each other via subsumption or sibling relationships (see Section 5.2). Comparisons of curator annotations from this experiment show that, on average, only 26% of character-state comparisons are exact matches. Given that the majority of curator annotation pairs are partial matches, the use of semantic similarity metrics that can quantify different degrees of similarity proves to be important.

### 5.5 Effect of external knowledge on inter-curator consistency and accuracy

One of the major differences between human and machine annotation is that humans can access external knowledge during curation, while machines cannot, beyond the encoded knowledge they have access to (here in the form of ontologies). Our measures of semantic similarity agreed with the results of Cui et al. (11) in showing that access to external knowledge had no effect on inter-curator consistency and did not further differentiate them from SCP’s annotations. Further, similarity to the Gold Standard was not generally increased. This was true despite the fact that curators changed annotations considerably between the Naïve and Knowledge rounds. Interestingly, while we expected a general increase in complexity when curators were at liberty to bring in additional knowledge, this was not borne out by the data.

These results indicate that lack of access to external knowledge is not one of the factors that contributes to SCP’s low performance with respect to human curators. This is encouraging, because lack of access to external knowledge during machine curation would be a challenge to remedy.

### 5.6 Machine performance is improved as ontologies become more complete

Our results indicate that using more complete ontologies can significantly improve machine performance (Figures 4 and 5). This is encouraging because ontology completeness is continually improved through the synergistic efforts of the ontology and curator communities.

This finding leads to specific ideas for how the curation workflow could be optimized by alternating execution of steps between human curators and algorithms. For instance, an initial round of machine curation would identify character states in the dataset for which good ontology matches were not found. Subsequently, human curators would judge whether the input ontology contains appropriate terms and focus on problem areas to add missing terms accordingly. Machines would then proceed with annotation using the human curator enhanced ontologies. Subsequently, human curators would review machine annotations and then either accept, modify, or re-curate them on a per-annotation basis. In such a workflow, machines would valuably augment the work of humans in the annotation process.

### 5.7 Future Work

#### 5.7.1 Improving reasoning over EQ annotations

One of the major challenges with EQ annotations is efficiently calculating semantic similarity metrics. Specifically, for virtually all metrics, the first step is to identify common subsuming classes. Although in theory an OWL reasoner can perform this task, it can only identify named classes that already exist in the ontology. A brute-force approach in which a composite ontology is computed as the cross-product of *E × Q × RE* terms (for entity, quality, related entity; or even only *E* × *Q*) (55) would result in a background ontology too large even for efficient reasoners such as ELK, and the vast majority of its compound classes would not be needed as subsumers. Further work is needed to improve this method for efficiency (computational time and memory) of the semantic similarity scoring.

#### 5.7.2 Improving Semantic CharaParser

Cui et al. (11) identified a number of areas of potential improvement for SCP, and the present study further refines our understanding of where the machine curation is encountering obstacles. The observed shortcomings primarily fall in the areas of entity post-composition, the handling of relational qualities in annotations, and ontology searching in PATO. One way to improve the latter would be to enable the ontology search to locate multiple-word PATO qualities such as *‘posteriorly directed’*, which in turn would allow more meaningful post-composed terms to be generated. And mentioned in Section 5.6, our results show that more comprehensive input ontologies will lead to improved performance of SCP.

## Conclusions

The Gold Standard dataset for EQ phenotype curation developed herein is a high-quality resource that will be of value to the sizable community of biocurators annotating phenotypes using the EQ formalism. As illustrated here, the Gold Standard enables assessment of how well a machine can performs EQ annotation and the impact of using different ontologies for that task. At present, machine-generated annotations are less similar to the Gold Standard than those of an expert human curator. The continued use of this corpus as a Gold Standard will enable training and evaluation of machine curation software in order to ultimately make phenotype annotation accurate at scale.

## Acknowledgments

This work was supported by the National Science Foundation (DBI-1062542, EF-0849982, and DBI-1147266). We thank M. Haendel and C. Mungall for suggestions on experiment design and ontology usage. We thank P. Chakrabarty, J. Clark, M. Coates, R. Hill, P. Skutschas, and J. Wible for completing the author assessments. P. Fernando and L. Jackson provided valuable feedback on the Gold Standard and Phenoscape Guide to Character Annotation. We thank D. Blackburn for his helpful comments on this manuscript. This manuscript is based on work done by P. Mabee while serving at the U.S. National Science Foundation. The views expressed in this paper do not necessarily reflect those of the National Science Foundation or the United States Government.

